# Control of stem cell behavior by CLE–JINGASA signaling in the shoot apical meristem in *Marchantia polymorpha*

**DOI:** 10.1101/2023.06.23.545176

**Authors:** Go Takahashi, Tomohiro Kiyosue, Yuki Hirakawa

**Affiliations:** Department of Life Science, Graduate School of Science, Gakushuin University, Tokyo 171-8588, Japan

**Keywords:** Marchantia, land plant, meristem, stem cell, CLE peptide, JINGASA, periclinal division

## Abstract

Land plants undergo indeterminate growth by the activity of meristems in both gametophyte (haploid) and sporophyte (diploid) generations^1-3^. In the sporophyte of the flowering plant *Arabidopsis thaliana*, the apical meristems are located at the shoot and root tips, in which a number of regulatory gene homologs are shared for their development, implying deep evolutionary origins^4-7^. However, little is known about their functional conservation with gametophytic meristems in distantly related land plants such as bryophytes^8-17^, even though genomic studies have revealed the subfamily-level diversity of regulatory genes is mostly conserved throughout land plants^18-23^. Here we show that a NAM/ATAF/CUC (NAC) domain transcription factor, JINGASA (MpJIN), acts downstream of CLV3/ESR-related (CLE) peptide signaling and controls stem cell behavior in the gametophytic shoot apical meristem of the liverwort *Marchantia polymorpha*. We found that Mp*JIN* shows heterogeneous expression in space and time within the stem cell zone, associated with the loss and *de novo* specification of stem cell identity at the branching event. Consistent with the expression pattern, induction of Mp*JIN* results in ectopic periclinal cell division in the stem cell zone and meristem termination. Comparative expression analysis suggests that the function of JIN/FEZ subfamily genes was shared between the shoot apical meristems in the gametophyte and sporophyte generations in early land plants but was lost in certain lineages including the flowering plant *A. thaliana*.

## Results

### MpCLE2 signaling negatively regulates the expression of *JINGASA*

In the apical notch of the *M. polymorpha* thalloid shoot, MpCLE2 signaling positively regulates the size of the meristematic cell population called the stem cell zone (SCZ)^14,24^. In contrast, the canonical CLV3 signaling pathway of flowering plants limits the stem cell population size via the transcription factor WUSCHEL^25,26^. Instead, the stem cell-promoting activity of MpCLE2 is suggested to be shared in *A. thaliana* CLE40 acting in the peripheral zone of the inflorescence meristem although its downstream signaling mechanisms are unknown^27,28^. To identify downstream targets of the stem cell-promoting MpCLE2 signaling, we performed transcriptome sequencing (RNA-seq) comparing wild type, Mp*CLE2* overexpression (OX) lines driven by the notch-specific Mp*YUCCA2* (Mp*YUC2*) promoter that exhibit an over-proliferation of stem cells, and knockout (KO) alleles for MpCLE2 receptor genes, Mp*CLV1* and Mp*CIK*^14,29^ (Fig. 1A and S1A). Compared to the wild-type transcriptome, hundreds of differentially expressed genes (DEGs) were found in each line, including less than 30 differentially expressed transcription factors (DETFs) (Fig. 1A and S1B). Among the latter, we found four DETFs specific to the MpCLE2-OX lines (Mp*NAC6*, Mp*NAC5*, Mp*ERF14*, and Mp*ASLBD11*) and one DETF specific to the receptor-KO lines (Mp*NAC7*) (Fig. 1B and S1C). Notably, Mp*NAC6* expression was dramatically decreased in the MpCLE2-OX lines (Fig. 1B and S1C). In qRT-PCR assays, the Mp*NAC6* mRNA level was decreased in the MpCLE2-OX line and was increased in an MpCLE2-KO allele^14^, *Mpcle2-3*^*ge*^ (Fig. 1C), suggesting that Mp*NAC6* expression is negatively regulated by MpCLE2 signaling. NAC proteins comprise of a large family of plant (streptophyte)-specific transcription factors, known to control various physiological processes including development, senescence, and stress responses^30^. Molecular phylogenetic analysis has classified MpNAC6 into a subfamily of the NAC family together with *A. thaliana* FEZ^21,30,31^ (Fig. S2A), and we designated MpNAC6 as JINGASA/MpJIN, inspired by the naming of FEZ which regulates root cap development in the *A. thaliana* root apical meristem^32^. Specific members in the JIN/FEZ subfamily were found in land plants including the moss *Sphagnum fallax* but not in the moss *Physcomitrium patens* or the hornwort *Anthoceros agrestis*^21,30^ (Fig. S2A), suggesting that JIN/FEZ subfamily has been established after gene duplications of NAC in the common ancestor of land plants and secondarily lost in some lineages.

**Figure 1.**
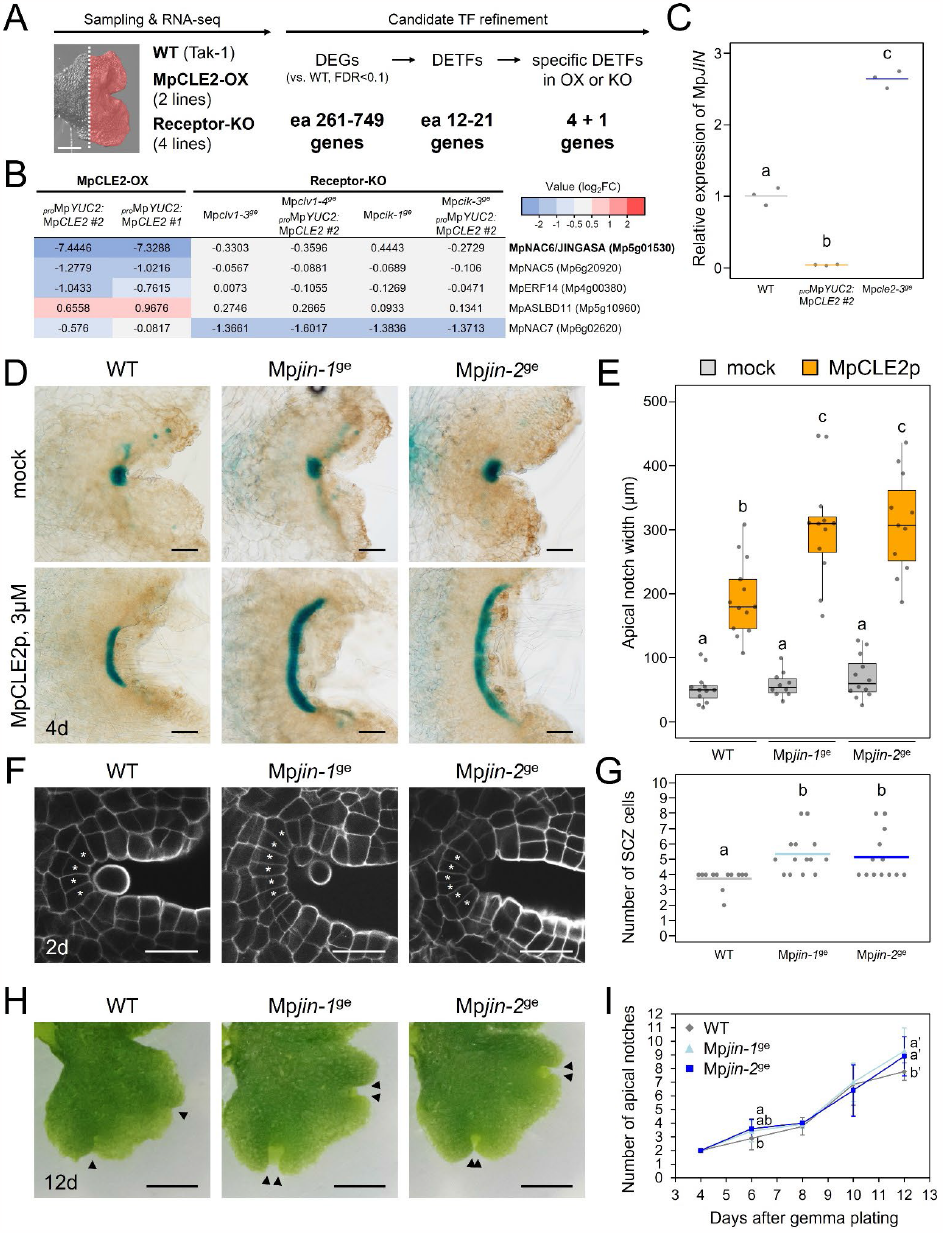
Mp*NAC6/JINGASA* negatively regulates the stem cell zone size under MpCLE2 signaling. (**A**) Schematic workflow of transcriptome analysis. (left) Sampled tissue (red) and genetic lines for RNA-seq. (right) Number of genes characterized as DEGs, DETFs and specific DETFs in OX lines or KO alleles (compared to WT, FDR<0.1). (**B**) Heat map illustration of the expression of the specific DETFs. Data represented in a log_2_ fold-change color code. (**C**) Relative expression levels of Mp*JIN* in 4-day-old gemmalings, normalized by Mp*APT* (n=3). (**D**) _*pro*_Mp*YUC2:GUS* marker in 4-day-old gemmalings grown with or without 3 μM MpCLE2 peptide. Genotypes of Mp*JIN* are indicated above the panels. (**E**) Quantification of the apical notch width (n=10-13). (**F**) Confocal imaging of the stem cell zone (SCZ) in 2-day-old gemmalings. Cell walls were stained with SCRI Renaissance 2200 (SR2200). Asterisks indicate the SCZ cells. (**G**) Quantification of the cell number in the SCZ (n=12-14). (**H**) Morphology of apical notch in 12-day-old gemmalings. Note that Mp*jin*^*ge*^ alleles show recently branched apical notches (arrowheads). (**I**) Quantification of the number of apical notches at 4, 6, 8, 10 and 12 days after gemmae plating (n=19-24). In **C** and **G**, data is represented by mean (bars) and individual data points (dots). In **E**, the boxes show the median and interquartile range, and the whiskers show the 1.5x interquartile range. Individual data points are plotted as dots. In **I**, the line plots show the mean with S.D. Two-way ANOVA with Tukey’s post hoc test in **C, E, G** and **I**. Means sharing the superscripts are not significantly different from each other, p < 0.05. Scale bars represent 500 μm (**A**), 100 μm (**D**), 25 μm (**F**), 250 μm (**H**).

### Genome editing of *JINGASA* affects the meristem activity

To elucidate the function of MpJIN in the shoot apical meristem of *M. polymorpha*, we generated loss-of-function alleles for Mp*JIN* using CRISPR-Cas9 editing^33^ in a _*pro*_Mp*YUC2:GUS* background, marking the apical notch^34^. Two independent frame-shift mutants (Mp*jin-1*^*ge*^ and Mp*jin-2*^*ge*^) at different target sites were obtained and used for further analysis (Fig. S2B and S2C). To examine their involvement in MpCLE2 signaling, we observed the phenotypes in gemmalings grown for 4 days on liquid medium supplemented with or without MpCLE2 peptides^14^. In the mock treatment, we could not find significant differences between Mp*jin-1*^*ge*^, Mp*jin-2*^*ge*^, and WT (in _*pro*_Mp*YUC2:GUS* background) in both apical notch morphology and GUS marker staining (Fig. 1D and 1E). However, when grown with 3 μM MpCLE2 peptides, Mp*jin-1*^*ge*^ and Mp*jin-2*^*ge*^ showed increased enlargement of the apical notch compared to WT, coinciding with wider _*pro*_Mp*YUC2:GUS* staining (Fig. 1D and 1E). These data indicate that the loss of Mp*JIN* function causes a hyper-responsive phenotype to exogenous MpCLE2 peptides. To characterize the function of Mp*JIN* in cellular resolution, we observed the SCZ of the apical notch using 2-day-old gemmalings grown in normal conditions (without peptide). In Mp*jin-1*^ge^ and Mp*jin-2*^ge^, cells in the SCZ were slightly increased compared to WT (Fig. 1F and 1G), which can be complemented by introducing the Mp*JIN* coding sequence expressed under its own promoter (Fig. S2D). These data show that Mp*JIN* negatively regulates the size of SCZ cell population. *M. polymorpha* thalloid shoots undergo periodic dichotomous branching by the duplications of apical notches. In time course analyses, Mp*jin-1*^*ge*^ and Mp*jin-2*^*ge*^ plants branched slightly faster compared to WT, implying that the increased SCZ population size may lead to more frequent branching (Fig. 1H and 1I).

### Expression dynamics of *JINGASA* in the stem cell zone

To understand the mechanism of action of MpJIN in the SCZ, we analyzed the spatial expression pattern of Mp*JIN* by generating transcriptional reporter fusion lines (_*pro*_Mp*JIN:3xCitrine*). In 2-day-old gemmalings of _*pro*_Mp*JIN:3xCitrine*, a fluorescence signal was detected broadly at the central region of apical notch (Fig. S2E). To observe the expression at the SCZ, we further performed confocal imaging with the iTOMEI clearing method^35^ (Fig. 2A and 2B). In the epidermal cell layer of the apical meristem including the SCZ, fluorescence signals were heterogeneous among cells: 1) the signals were faint at the center of the SCZ, 2) slightly higher signals were detected in adjacent cells in the SCZ, and 3) the signals were at the maxima at the periphery of the SCZ where two cell layers (epidermal and subepidermal layers) were formed after a recent periclinal cell division, judged from the cellular arrangement. This pattern implies that the strong Mp*JIN* expression is associated with the periclinal division. Furthermore, cells with faint Mp*JIN* expression (MpJIN-negative cells) within the SCZ could distinguish the apical cell from its immediate derivatives (subapical cells) with similar morphology^36-39^. In the MpCLE2-OX lines (_*pro*_Mp*YUC2:*Mp*CLE2*), the SCZ is expanded broader than wild type, from which multiple meristems/branches will arise^14^. In this genetic background, MpJIN-negative and MpJIN-positive cells were alternately aligned in the expanded SCZ at day 2, forming multiple MpJIN-negative spots within the SCZ (Fig. 2C and 2D). Obvious morphological differences were not observed between the cells in the SCZ at this time point, suggesting that the heterogeneity of Mp*JIN* expression precedes the morphological differentiation of the SCZ cell population at the branching event.

**Figure 2.**
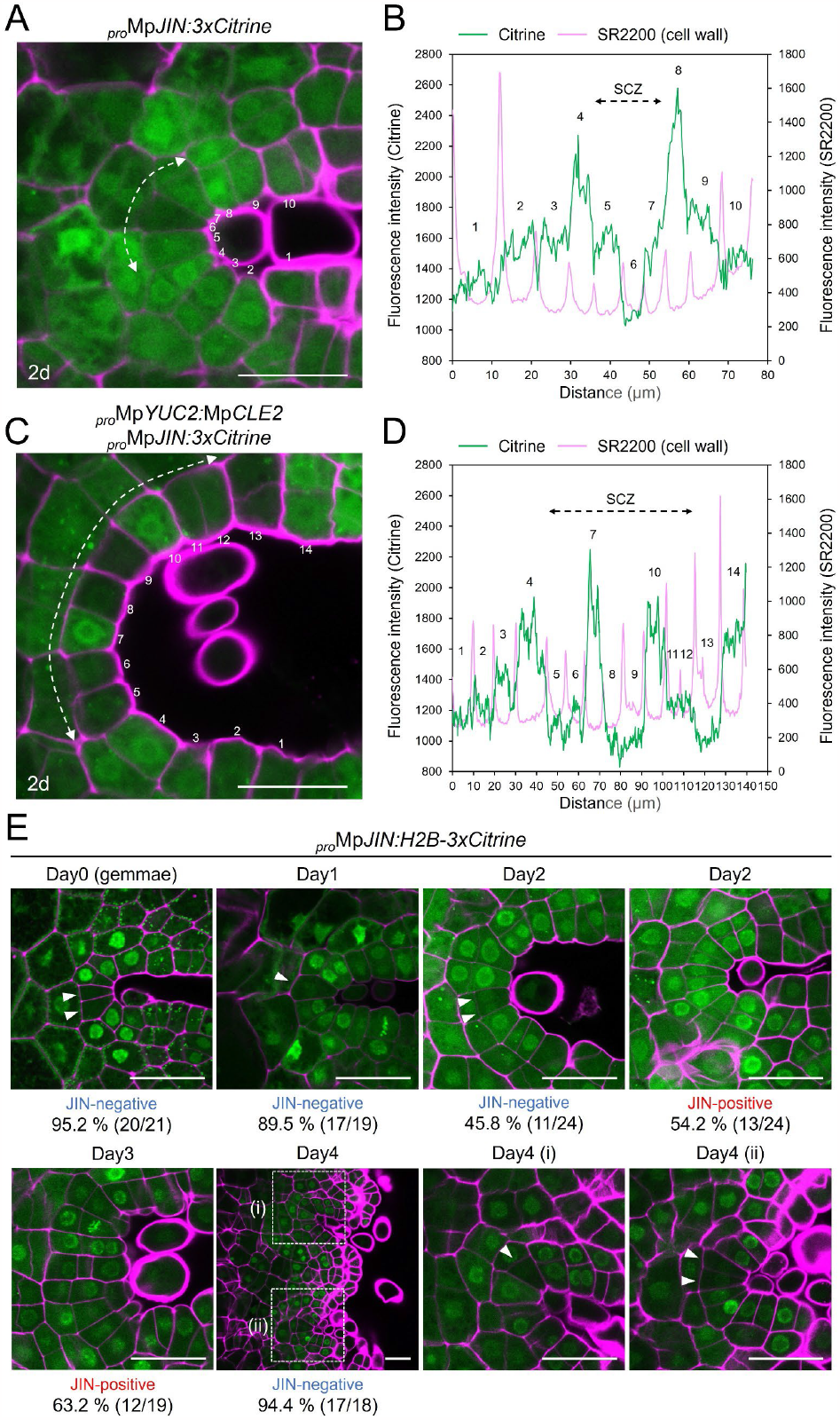
Spatio-temporal patterns of Mp*JIN* promoter activity in the SCZ. (**A**-**D**) Confocal imaging of _*pro*_Mp*JIN:3xCitrine* in the SCZ in 2-day-old gemmaling in WT (**A**,**B**) or MpCLE2-OX (**C**,**D**). Cell walls were stained with SCRI Renaissance 2200 (SR2200). Dashed lines indicate the SCZ. Numbers indicate the position of cells in fluorescence intensity measurements. (**E**) Time-course confocal imaging of _*pro*_Mp*JIN:H2B-3xCitrine* during branching. Arrowheads indicate JIN-negative cells in the SCZ. Days after gemmae plating are indicated above the panels. The ratios of SCZs containing at least one JIN-negative cell (JIN-negative) or none (JIN-positive) are indicated below the panels (n=18-24). Day 4 (i) and (ii) panels are the magnified images of the dashed line squares in Day 4 panel. Scale bars represent 25 μm (**A**,**C**,**E**).

To analyze the time course of Mp*JIN* promoter activity in the normal branching event in gemmaling development, we generated a nuclear-localized reporter (_*pro*_Mp*JIN:H2B-3xCitrine*) for enhanced signal intensity. In _*pro*_Mp*JIN:H2B-3xCitrine*, JIN-negative cells were often observed as a group of 1 to 6 cells in the SCZs in gemmae (day 0), which were decreased to 1 or 2 cells in most gemmalings at day 1. We measured the ratio of SCZs containing at least one JIN-negative cell (JIN-negative SCZ) during gemmaling development (n=18-24 at each time point, Fig. 2E). At day 0 (gemmae), the ratio of JIN-negative SCZ was 95.2% of observations, and a similar ratio of JIN-negative SCZs (89.5%) was observed at day 1. At day 2, however, the ratio of JIN-negative SCZ decreased to 45.8%. Instead, 54.2% SCZs showed relatively homogenous Mp*JIN* expression in the SCZ, suggesting upregulation of Mp*JIN* expression (MpJIN-positive SCZ). Similarly, MpJIN-positive SCZs were found in 63.2% of observations at day 3. During day 3 to day 4, new SCZs arise from the two ends of the original SCZ, separated by a central protruding tissue (central lobe). At day 4, JIN-negative SCZs were found at one or two ends of the original SCZ in 94.4% of observation (Fig. 2E). Collectively, these data suggest that the JIN-negative stem cell identity is lost from the SCZ before the branching event and respecified *de novo* at two separate positions.

### Induction of periclinal division by *JINGASA* in the stem cell zone

To further investigate the cellular function of MpJIN, we generated inducible overexpression lines in which an MpJIN-glucocorticoid receptor fusion protein (MpJIN-GR) is expressed under the constitutive Mp*EF1α* promoter^40^. By DEX treatment, MpJIN-GR is expected to be translocated into nucleus and affect the transcription of target genes. In DEX-containing medium, _*pro*_Mp*EF1α:*Mp*JIN-GR* plants showed a significant reduction of overall growth in a dose-dependent manner (Fig. 3A and 3B). In the apical notches at day 2, ectopic periclinal cell divisions occurred in a cuneate-shaped stem cell at the center of the SCZ in 22.8 % of observations (n=19, Fig. 3C), which was never observed in DEX-free condition (n=18, Fig. 3C). To analyze the influence of the ectopic periclinal cell division on meristem maintenance, _*pro*_Mp*EF1α:*Mp*JIN-GR* plants were transferred from DEX-containing medium to DEX-free medium at day 3. Three days after the transfer, the _*pro*_Mp*YUC2:GUS* marker was not detected at the original apical notch (Fig. 3D), suggesting the loss of the meristem identity. Instead, GUS-positive spots with mucilage papillae appeared in small notches at the periphery of the thallus (Fig. 3D), which may have happened as a regeneration process after the loss of original apical meristem activity. Collectively, these data suggest that ectopic expression of Mp*JIN* in the SCZ induces periclinal division and causes the loss of stem cell activity.

**Figure 3.**
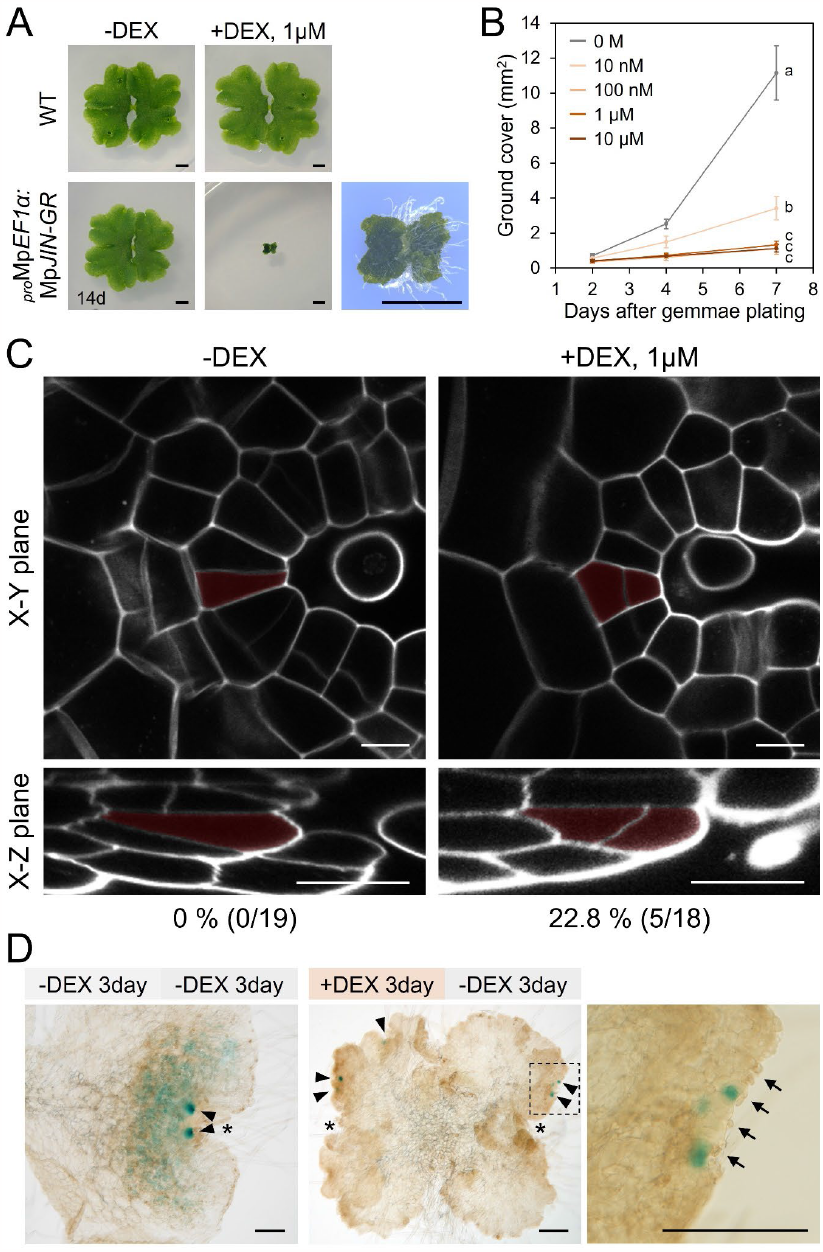
Overexpression of Mp*JIN* induces periclinal division in the SCZ and meristem termination. (**A**) Overall morphology of 14-day-old plants grown from gemmae with or without 1 μM dexamethasone (DEX). The lower right panel shows the magnified image of _*pro*_Mp*EF1α:*Mp*JIN-GR* plant grown with DEX. (**B**) Quantification of the ground cover area of _*pro*_Mp*EF1α:*Mp*JIN-GR* plants grown with different DEX concentrations at 2, 4 and 7 days after gemmae plating (mean and SD, n=12). (**C**) Confocal imaging of the SCZ in 2-day-old _*pro*_Mp*EF1α:*Mp*JIN-GR* plants with or without 1 μM DEX. Cell walls were stained with SCRI Renaissance 2200 (SR2200). SCZ cells observed in X-Y and X-Z (z-stack) planes are coloured red. The ratios of the SCZs with ectopic periclinal division are indicated below (n=18-19). (**D**) Effects of temporal Mp*JIN* induction on apical notch maintenance. _*pro*_Mp*EF1α:*Mp*JIN-GR* gemmae were grown for 3 days on DEX-free or DEX-containing medium and then transferred to DEX-free medium for additional 3-day culture. The right panel is the magnified image of dashed line square in the middle panel. Asterisks, arrowheads and arrows indicate original apical notches, _*pro*_Mp*YUC2:GUS* spots and mucilage cells, respectively. Two-way ANOVA with Tukey’s post hoc test in **B**. Means sharing the superscripts are not significantly different from each other, p < 0.05. Scale bars represent 250 μm (**A**), 10 μm (**C**), 200 μm (**D**).

## Discussion

Based on these results, we propose that a NAC-domain protein, MpJIN, regulates stem cell behavior in the shoot apical meristem of *M. polymorpha* under the control of MpCLE2 peptide signaling. High-level expression of Mp*JIN* promotes periclinal cell division at the lateral periphery of the SCZ, which results in the formation of two daughter cells at the epidermal and subepidermal positions (Fig. 4A). Mp*JIN* has a negative impact on the maintenance of stem cell identity because the loss of Mp*JIN* function results in the increased SCZ size and faster branching (Fig. 1D-1I), and the ectopic Mp*JIN* activation results in the meristem termination (Fig. 3D). The stem cell-limiting function of Mp*JIN* is consistent with the stem cell-promoting function of MpCLE2^14^ since MpCLE2 signaling reduces the expression of Mp*JIN* (Fig. 4B). However, the meristem expansion phenotypes of Mp*jin*^*ge*^ alleles were mild compared to MpCLE2-OX lines^14^. Furthermore, Mp*jin*^*ge*^ alleles were (hyper)sensitive to MpCLE2 peptide, indicating that MpCLE2 signaling must also have an unknown target gene(s) other than Mp*JIN* (Fig. 4B). Our transcriptome analysis provides additional DETFs as potential targets of MpCLE2 signaling. One of which, Mp*NAC5* (Mp6g20920), belongs to the VND/NST/SMB (VNS) subfamily of the NAC protein family together with *A. thaliana SOMBRERO* (*SMB*), known to form a feedback regulatory loop with *FEZ* in the root cap stem cell maintenance^21,32,41^. However, CRISPR editing of Mp*NAC5* (Mp*nac5-1*^*ge*^/*-2*^*ge*^) did not cause significant changes in either the responsiveness to MpCLE2 peptide or the SCZ size (Fig. S3), suggesting that Mp*NAC5*/Mp*VNS* is not important for the regulation of stem cell dynamics, unlike the case of *SMB* in the *A. thaliana* root cap stem cells^32^.

**Figure 4.**
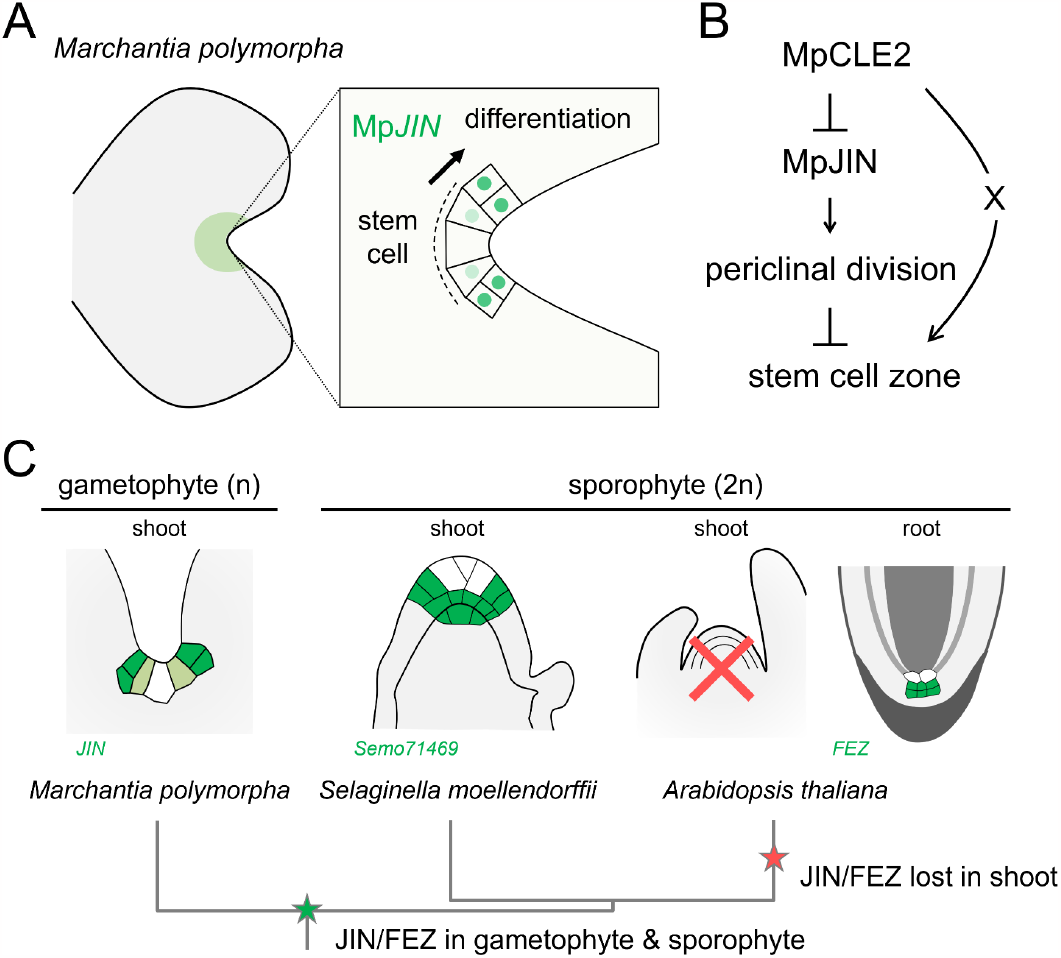
Models for the function and evolution of JIN/FEZ subfamily in land plants. (**A**) Model for Mp*JIN* function in the control of stem cell behavior in *M. polymorpha*. (**B**) Model for MpCLE2 signaling pathway mediated by MpJIN and unknown factor ‘X’. (**C**) Model for the role of JIN/FEZ subfamily in the evolution of apical meristems in land plants. Phylogenetic relationships of species are indicated below with stars representing the timing of gain and loss of JIN/FEZ function. Expression of JIN/FEZ near the stem cells are illustrated (green, expressed; white, not expressed).

Our results show that Mp*JIN* acts in the shoot apical meristem of *M. polymorpha*. In contrast, the expression and function of *A. thaliana* homologs, *FEZ* and *FEZ-LIKE1*, have been reported only in the root apical meristem^32,42^, suggesting that the function of *JIN/FEZ* is not conserved between the shoot apical meristems of these species (Fig. 4C). However, we found that a *JIN/FEZ* homolog (*Semo71469*) of the lycophyte *Selaginella moellendorffii* is preferentially expressed at the ‘meristem core’ that surrounds the apical cell within the sporophytic (diploid) shoot apical meristem in gene expression atlas^43,44^ (Fig. S4), showing similarity with the Mp*JIN* expression pattern in *M. polymorpha* gametophyte (Fig. 4C). It is noteworthy that some vascular plants, such as lycophytes and *Psilotum nudum*, undergo dichotomous branching, which is regarded as an ancestral characteristic of growth from the meristem. Anatomical studies of *P. nudum* aerial shoots have shown that the original apical cell is inactivated, and two apical cells are formed *de novo* during dichotomous branching^45^, which has a similarity to the Mp*JIN* expression patterns in *M. polymorpha*, although the mode of branching is variable between species and tissues^46^. Based on these findings, we propose that the control of stem cell division by *JIN/FEZ* is a mechanism shared between the gametophytic and sporophytic shoot apical meristems in early land plants and is preserved in the shoot apical meristems of some extant bryophytes and vascular plants (Fig. 4C). The subfamily-level diversity in NAC family, including JIN/FEZ subfamily, is estimated to be established in the common ancestor of land plants^21,23,30^. In the 2D-structured body of *Coleochaete* algae, the position of periclinal cell divisions can be explained by simple, cell shape-based rules^47^. It is conceivable that in the common ancestor of land plants a JIN/FEZ subfamily protein evolved that allowed precise spatio-temporal regulation of periclinal divisions in the stem cell population. Similarly, some GRAS family transcription factors are reported to promote or inhibit periclinal divisions in *A. thaliana* and the moss *P. patens*^48^. The expansion of gene families by gene duplications likely has contributed to the evolution of complexity in the plant body. The *JIN/FEZ* gene expression in the root cap is shared between *S. moellendorffii* (*Semo25673*) and *A. thaliana*^49^ (Fig. S4). The two plant lineages are estimated to have split early in vascular plant evolution and have innovated the roots independently, suggesting independent recruitments of *JIN/FEZ* in the evolution of root apical meristems^49^. In this scenario, the function of JIN/FEZ must have been lost in the shoot apical meristem in the flowering plant *A. thaliana* (Fig. 4C), in which periclinal cell divisions are largely excluded to maintain its layered tissue structure called tunica^50,51^. Taken together, our findings illuminate complicated evolutionary paths toward the diverse meristem structures in extant land plants, which may have hindered the identification of conserved regulatory mechanisms in previous studies using a small number of model plants. Future studies with broader taxon sampling will improve our fundamental understanding of the development and evolution of plant meristems.

## Methods

### Plant materials and growth conditions

*Marchantia polymorpha* male Takaragaike-1 (Tak-1) accession was used as wild type in this study. *M. polymorpha* plants were grown on half-strength Gamborg B5 medium (pH 5.5) solidified with 1.4% agar at 22°C under continuous white light.

### Plasmid Construction

Primers used in this study are listed in Table S1. All plant transformation vectors were generated using the Gateway cloning system (Thermo Fisher Scientific, Waltham, MA, United States). Gateway destination vectors are described in Ishizaki et al. (2015)^52^ and Sugano et al. (2018)^33^.

To generate H2B-3×Citrine Gateway binary vectors, the coding sequence of a histone 2B gene (At1g07790/HTB1) was PCR amplified from *Arabidopsis thaliana* genomic DNA with a primer pair of AtH2B_CDS_F_Aor51HI and AtH2B_CDS_R_Aor51HI, and inserted into the *Aor*51HI-linearized pMpGWB123/223/323/423 vectors using In-Fusion HD Cloning Kit (Takara Bio, Shiga, Japan).

For promoter reporter analysis, a 4609 bp DNA fragment of Mp*JIN* promoter sequence flanking the translational initiation site was PCR amplified with a primer pair of MpJINprom4.6k_F and MpJINprom_R, and cloned into pENTR/D-TOPO vector (Thermo Fisher Scientific). The resulting plasmid, pENTR-MpJINp, was transferred to the pMpGWB323 or pMpGWB323-H2B vector using Gateway LR Clonase II Enzyme mix (Thermo Fisher Scientific).

For genome editing of Mp*JIN* and Mp*NAC5*, guide RNA was designed at the coding sequence of Mp5g01530 and Mp6g20920, respectively, using CRISPRdirect^53^. The plasmids for genome editing were constructed according to Sugano et al. (2018)^33^.

For production of inducible Mp*JIN* overexpression alleles, the coding sequence of Mp*JIN* was PCR amplified from *M. polymorpha* cDNA with a primer pair of MpJINCDS_F and MpJINCDS_R_-stop, and cloned into pENTR/D-TOPO vector. The resulting vector, pENTR-MpJINcds_-stop, was transferred to pMpGWB113 using Gateway LR Clonase II Enzyme mix (Thermo Fisher Scientific).

For complementation of genome editing alleles for Mp*JIN*, the 4609 bp Mp*JIN* promoter sequence was PCR amplified with a primer pair of MpJINprom4.6k_F_InFusion_XbaI and MpJINprom_R_InFusion_XbaI, and cloned into the *Xba*I digestion site of pMpGWB307 vectors using In-Fusion HD Cloning Kit (Takara Bio) to produce pMpGWB307-MpJINp. The coding sequence of Mp*JIN* was PCR amplified from *M. polymorpha* cDNA with a primer pair of MpJINCDS_F and MpJINCDS_R_+stop, and cloned into pENTR/D-TOPO vector. pENTR-MpJINcds_+stop was transferred to pMpGWB307-MpJINp using Gateway LR Clonase II Enzyme mix (Thermo Fisher Scientific).

### Transgenic plants

Transgenic *M. polymorpha* plants are listed in Table S2. *Agrobacterium*-mediated transformation of *M. polymorpha* was performed using regenerating thalli according to Kubota et al. (2013)^54^. CRISPR/Cas9-based genome editing was performed according to Sugano et al. (2018)^33^. Mutations in the guide RNA target loci were examined by direct sequencing of PCR product amplified from genome DNA samples with primers listed in Table S1.

### Peptide treatment

Synthetic peptides used in this study were analytically pure and dissolved in 0.1% TFA (trifluoroacetic acid) solution as stock solutions. For MpCLE2 peptide treatment, approximately 20 mature gemmae were floated on 2 mL liquid M51C medium containing 2% sucrose supplemented with 3 μM MpCLE2 peptide (KEVHypNGHypNPLHN) or mock (TFA) solution, in 12-well plates as described previously^14^.

### GUS staining

Individual plants were stained separately in 30–50 uL GUS staining solution (50 mM sodium phosphate buffer pH 7.2, 1 mM potassium-ferrocyanide, 1 mM potassium-ferricyanide, 10 mM EDTA, 0.01% Triton X-100 and 1 mM 5-bromo-4-chloro-3-indolyl-b-D-glucuronic acid) at 37 C in dark. GUS-stained samples were washed with water, cleared with ethanol, and mounted with clearing solution (chloral hydrate-glycerol-water, 8:1:2) for imaging under a light microscope (BX51, Olympus, Tokyo, Japan) as described previously^14^.

### Imaging and phenotypic measurement

Measurements were taken from distinct samples except for those in Fig. 1e and 3b where the same sample was measured repeatedly. Overall morphology of plants and the measurement of apical notch number, plants were imaged under a digital microscope (DMS1000, Leica Microsystems, Wetzlar, Germany) or using a digital camera (TG-6, Olympus). To quantify the apical notch width, distance between the rims of apical notch was measured on the images using ImageJ^55^. For measurement of the SCZ size, 2-day-old gemmalings were fixed and cleared with ClearSee protocol^56^ as described previously^14^. For florescent observation, plants were fixed and cleared with iTOMEI protocol^35^. Briefly, plants were fixed with 1% paraformaldehyde in PBS solution for 1 hour with evacuation for the first 10 minutes (0.06 MPa). After fixation, the plants were cleared for overnight with gentle shaking in decolorization solution (20% caprylyl sulfobetaine [TCI, Tokyo, Japan] in 100 mM sodium phosphate buffer at pH 8.0). Cleared plants were stained for 1 hour with 1% SCRI Renaissance 2200 (Renaissance Chemicals, Selby, United Kingdom) in PBS solution, and then incubated in the mounting solution (70.4% iohexol [TCI, Japan] in PBS solution) for 1 hour with gentle shaking. For the observation of the SCZ at the ventral side of meristem, gemmalings were fixed and cleared with small portion of agar medium attached via rhizoids. Finally, samples were mounted in the mounting solution and observed under a confocal laser scanning microscopy (Fluoview FV1000, Olympus or Fluoview FV3000, Olympus). Z-series images were collected at 0.5 μm intervals through the specimens and obtained images were processed using ‘Crop(3D)’ function of Fiji software^55^ to specify the SCZ. Fluorescent intensities were measured at arbitrary lines which are parallel to the epidermal surface and cross the center of epidermal cells, using ‘Plot profile’ function of Fiji software.

### qRT-PCR

To quantify Mp*JIN* mRNA level, total RNA was extracted from tissues including apical notches in 4-day-old gemmalings using NucleoSpin RNA Plant (Macherey-Nagel, Duren, Germany) according to manufacturer’s instruction, and first-strand cDNAs were prepared using ReverTra Ace qPCR RT Master Mix with gDNA Remover (TOYOBO, Osaka, Japan). qRT-PCR was performed TB Green Premix Ex Taq II (Takara) and the StepOnePlus Real-Time PCR System (Thermo Fisher Scientific). Mp*APT* (Mp3g25140) was used as reference gene^57^. To prevent amplification from genomic DNA, primer pairs were designed at exon/intron junction as listed in Table S1.

### Transcriptome analysis

For transcriptome analysis, total RNA was extracted from thallus of 4-day-old gemmalings including apical notches, using NucleoSpin RNA Plant (Macherey-Nagel). The quality and concentration of total RNA was measured by Agilent 2100 bioanalyzer (Agilent, Santa Clara, CA, United States).

Sequence libraries were prepared using a TruSeq Stranded mRNA Sample Prep Kit (Illumina, San Diego, CA, United States) and sequenced using Ilumina Hiseq 2500 with a single-end sequencing protocol. Sequence data were deposited in the DDBJ Sequence Read Archive (DRA016127). The quality of sequence reads was checked by FastQC (https://www.bioinformatics.babraham.ac.uk/projects/fastqc/). After removing adaptor sequences by TrimGalore-0.6.5 with the following parameters: ‘--clip_R1 11 -- three_prime_clip_R1 3’, the sequence reads were mapped to the *M. polymorpha* reference genome MpTak1v5.1^31^ by STAR-2.7.4a^58^ with the following parameters: ‘--quantMode TranscriptomeSAM’. The mapped reads were counted using the RSEM-1.3.3^59^.

RNA samples were classified by PCA using iDEP.96 (integrated Differential Expression and Pathway analysis, http://bioinformatics.sdstate.edu/idep96/). Log2 fold changes and adjusted P-values (FDR) of mutant data were calculated by comparison with wild-type data. The TCC-GUI package (https://infinityloop.shinyapps.io/TCC-GUI/) was used to detect differentially expressed genes (DEGs) (FDR<0.1). Differentially expressed transcription factors (DETFs) were selected based on the gene list of transcription factors in *M. polymorpha*^21^.

### Phylogenetic analysis

Protein sequences were retrieved from the following databases: MarpolBase (https://marchantia.info), Phytozome (https://phytozome-next.jgi.doe.gov/)^60^, TAIR (http://www.arabidopsis.org/), ONEKP (https://db.cngb.org/onekp/)^61^ and GinkgoDB (https://ginkgo.zju.edu.cn/genome/)^62^. Alignment was performed on the amino acid sequences of the NAC domain using CLUSTALW (https://www.genome.jp/tools-bin/clustalw). After manually removing the alignment gaps using SeaView (https://doua.prabi.fr/software/seaview), phylogenetic analysis was performed on the alignment using MrBayes3.2.7^63^. Two runs with four chains of Markov chain Monte Carlo (MCMC) iterations were performed for 300,000 generations, keeping one tree every 100 generations. The first 25% of the generations were discarded as burn-in and the remaining trees were used to calculate a 50% majority-rule tree. The standard deviation for the two MCMC iteration runs was below 0.01, suggesting that it was sufficient for the convergence of the two runs. Convergence was assessed by visual inspection of the plot of the log likelihood scores of the two runs calculated by MrBayes^64^. Character matrix used to run the Bayesian phylogenetic analysis is provided in Supplementary Data 1.

### Data visualization

PlotsOfData^65^ (https://huygens.science.uva.nl/PlotsOfData/) was used for data visualization.

## Acknowledgments

We thank our colleagues at the Platform for Advanced Genome Science (PAGS) project for transcriptome analysis; Takashi Ueda, Takehiko Kanazawa, and Ikuko Nakanomyo for technical assistance; John L. Bowman, and John P. Alvarez for providing feedback on the manuscript. This work was supported by JSPS KAKENHI Grant Number JP19K06727, JP22H02676, and JP16H06279 (PAGS) to Y.H., and by NIBB Collaborative Research Program 21-241 to Y.H.

## Author contributions

Y.H. conceived the project. G.T. and Y.H. designed the experiments. G.T. performed the experiments with assistance from Y.H. G.T. and Y.H. analyzed the data, generated figures, and wrote the manuscript draft. All authors contributed to the article.

## References

1. Harrison, J.C. (2017) Development and genetics in the evolution of land plant body plans. Philos Trans R Soc Lond B Biol Sci 372, 20150490

2. Kenrick, P. (2018) Changing expressions: a hypothesis for the origin of the vascular plant life cycle. Philos Trans R Soc Lond B Biol Sci 373, 20170149

3. Bowman, J.L. et al. (2019) Evolution and co-option of developmental regulatory networks in early land plants. Curr Top Dev Biol 131, 35–53

4. Sarkar, A.K. et al. (2007) Conserved factors regulate signalling in Arabidopsis thaliana shoot and root stem cell organizers. Nature 446, 811–814

5. Barton, M.K. (2010) Twenty years on: the inner workings of the shoot apical meristem, a developmental dynamo. Dev Biol 341, 95–113

6. Engstrom, E.M. et al. (2011) Arabidopsis Homologs of the Petunia HAIRY MERISTEM Gene Are Required for Maintenance of Shoot and Root Indeterminacy. Plant Physiol 155:735–750

7. Scheres B., and Krizek B.A. (2018) Coordination of growth in root and shoot apices by AIL/PLT transcription factors. Curr Opin Plant Biol 41, 95–101

8. Graham, L.E. et al. (2000) The origin of plants: body plan changes contributing to a major evolutionary radiation. Proc Natl Acad Sci USA 97, 4535–4540

9. Aoyama, T. et al. (2012) AP2-type transcription factors determine stem cell identity in the moss Physcomitrella patens. Development 139, 3120–3129

10. Frank, M.H. & Scanlon, M.J. (2015) Transcriptomic evidence for the evolution of shoot meristem function in sporophyte-dominant land plants through concerted selection of ancestral gametophytic and sporophytic genetic programs. Mol Biol Evol 32, 355–367

11. Yip, K. et al. (2016) Class III HD-Zip activity coordinates leaf development in Physcomitrella patens. Dev Biol 419, 184–197

12. Whitewoods, C.D. et al. (2018) CLAVATA Was a Genetic Novelty for the Morphological Innovation of 3D Growth in Land Plants. Curr Biol 28, 2365–2376

13. Naramoto, S. et al. (2019) A conserved regulatory mechanism mediates the convergent evolution of plant shoot lateral organs. PLoS Biol 17, e3000560

14. Hirakawa, Y. et al. (2020) Induction of Multichotomous Branching by CLAVATA Peptide in Marchantia polymorpha. Curr Biol 30, 3833–3840

15. Hata, Y. & Kyozuka, J. (2021) Fundamental mechanisms of the stem cell regulation in land plants: lesson from shoot apical cells in bryophytes. Plant Mol Biol 107, 213–225

16. Beheshti, H, et al. (2021) PpGRAS12 acts as a positive regulator of meristem formation in Physcomitrium patens. Plant Mol Biol 107, 293–305

17. Fouracre, J.P. & Harrison, C.J. (2022) How was apical growth regulated in the ancestral land plant? Insights from the development of non-seed plants. Plant Physiol 190, 100–112

18. Nishiyama, T. et al. (2004) Chloroplast phylogeny indicates that bryophytes are monophyletic. Mol Biol Evol 21, 1813–1819

19. Rensing, S.A. et al. (2008) The Physcomitrella genome reveals evolutionary insights into the conquest of land by plants. Science 319, 64–69

20. Catarino, B. et al. (2016) The Stepwise Increase in the Number of Transcription Factor Families in the Precambrian Predated the Diversification of Plants On Land. Mol Biol Evol 33, 2815–2819

21. Bowman, J.L. et al. (2017) Insights into Land Plant Evolution Garnered from the Marchantia polymorpha Genome. Cell 171, 287–304

22. Wilhelmsson, P.K.I. et al. (2017) Comprehensive Genome-Wide Classification Reveals That Many Plant-Specific Transcription Factors Evolved in Streptophyte Algae. Genome Biol Evol 9, 3384–3397

23. Bowman, J.L. (2022) The origin of a land flora. Nat Plants 8, 1352–1369

24. Kohchi, T. et al. (2021) Development and Molecular Genetics of Marchantia polymorpha. Annu Rev Plant Biol 72, 677–702

25. Brand, U. et al. (2000) Dependence of stem cell fate in Arabidopsis on a feedback loop regulated by CLV3 activity. Science 289, 617–619

26. Schoof, H. et al. (2000) The stem cell population of Arabidopsis shoot meristems in maintained by a regulatory loop between the CLAVATA and WUSCHEL genes. Cell 100, 635–644

27. Schlegel, J. et al. (2021) Control of Arabidopsis shoot stem cell homeostasis by two antagonistic CLE peptide signalling pathways. Elife 10, e70934

28. Hirakawa, Y. (2022) Evolution of meristem zonation by CLE gene duplication in land plants. Nat Plants 8, 735–740

29. Takahashi, G. et al. (2021) An Evolutionarily Conserved Coreceptor Gene Is Essential for CLAVATA Signaling in Marchantia polymorpha. Front Plant Sci 12, 657548

30. Maugarny-Calès, A. et al. (2016) Apparition of the NAC Transcription Factors Predates the Emergence of Land Plants. Mol Plant 9, 1345–1348

31. Montgomery, S.A. et al. (2020) Chromatin Organization in Early Land Plants Reveals an Ancestral Association between H3K27me3, Transposons, and Constitutive Heterochromatin. Curr Biol 30, 573–588

32. Willemsen, V. et al. (2008) The NAC domain transcription factors FEZ and SOMBRERO control the orientation of cell division plane in Arabidopsis root stem cells. Dev Cell 15, 913–922

33. Sugano, S.S. et al. (2018) Efficient CRISPR/Cas9-based genome editing and its application to conditional genetic analysis in Marchantia polymorpha. PLoS One 13, e0205117

34. Eklund, D.M. et al. (2015) Auxin Produced by the Indole-3-Pyruvic Acid Pathway Regulates Development and Gemmae Dormancy in the Liverwort Marchantia polymorpha. Plant Cell 27,1650–1669

35. Sakamoto, Y. et al. (2022) Improved clearing method contributes to deep imaging of plant organs. Commun Biol 5, 12

36. Douin, R. (1921) Reserches sur les Marchantiées. Rev Gen Bot 33, 34–62; 99–145;190–209

37. Apostolakos, P. et al. (1982) Studies on the Development of the Air Pores and Air Chambers of Marchantia paleacea: 1. Light Microscopy. Ann Bot 49, 377–396

38. Shimamura, M. (2016) Marchantia polymorpha: Taxonomy, Phylogeny and Morphology of a Model System. Plant Cell Physiol 57, 230–256

39. Suzuki, H. et al. (2020) Positional cues regulate dorsal organ formation in the liverwort Marchantia polymorpha. J Plant Res 133, 311–321

40. Althoff, F. et al. (2014). Comparison of the MpEF1α and CaMV35 promoters for application in Marchantia polymorpha overexpression studies. Transgenic Res 23, 235–244.

41. Xu, B. et al. (2014) Contribution of NAC transcription factors to plant adaptation to land. Science 343, 1505–1508

42. Bennett, T. et al. (2014) Precise control of plant stem cell activity through parallel regulatory inputs. Development 141, 4055–4064

43. Frank, M.H. et al. (2015) Dissecting the molecular signatures of apical cell-type shoot meristems from two ancient land plant lineages. New Phytol 207, 893–904

44. Ferrari, C. et al. (2020) Expression Atlas of Selaginella moellendorffii Provides Insights into the Evolution of Vasculature, Secondary Metabolism, and Roots. Plant Cell 32, 853–870

45. Takiguchi, Y. et al. (1997) Cell division patterns in the apices of subterranean axis and aerial shoot of Psilotum nudum (Psilotaceae): morphological and phylogenetic implications for the subterranean axis. Am J Bot 84, 588

46. Gola, E.M. (2014) Dichotomous branching: the plant form and integrity upon the apical meristem bifurcation. Front Plant Sci 5, 263

47. Dupuy, L. et al. (2010) Coordination of plant cell division and expansion in a simple morphogenetic system. Proc Natl Acad Sci USA 107, 2711–2716

48. Ishikawa, M. et al. (2023) GRAS transcription factors regulate cell division planes in moss overriding the default rule. Proc Natl Acad Sci USA 120, e2210632120

49. Huang, L. & Schiefelbein, J. (2015) Conserved Gene Expression Programs in Developing Roots from Diverse Plants. Plant Cell 27, 2119–2132

50. Steeves, T.A. & Sussex, I.M. (1989) Patterns in plant development. Cambridge University Press

51. Evert, R.F. (2006) Esau’s Plant Anatomy: Meristems, Cells, and Tissues of the Plant Body: Their Structure, Function, and Development, 3rd Edition. Wiley-Liss

52. Ishizaki, K. et al. (2015) Development of Gateway Binary Vector Series with Four Different Selection Markers for the Liverwort Marchantia polymorpha. PLoS One 10, e0138876

53. Naito, Y. et al. (2015) CRISPRdirect: software for designing CRISPR/Cas guide RNA with reduced off-target sites. Bioinformatics 31, 1120–1123.

54. Kubota, A. et al. (2013) Efficient Agrobacterium-mediated transformation of the liverwort Marchantia polymorpha using regenerating thalli. Biosci Biotechnol Biochem 77, 167–172

55. Schneider, C.A. (2012). NIH Image to ImageJ: 25 years of image analysis. Nat Methods 9, 671–675

56. Kurihara, D. et al. (2015) ClearSee: a rapid optical clearing reagent for whole-plant fluorescence imaging. Development 142, 4168–4179

57. Saint-Marcoux, D. et al. (2015) Identification of reference genes for real-time quantitative PCR experiments in the liverwort Marchantia polymorpha. PLoS One 10, e0118678

58. Dobin, A. et al. (2013) STAR: ultrafast universal RNA-seq aligner. Bioinformatics 29, 15–21

59. Li, B. & Dewey, C.N. (2011) RSEM: accurate transcript quantification from RNA-Seq data with or without a reference genome. BMC Bioinformatics 12, 323

60. Goodstein, D.M. et al. (2012) Phytozome: a comparative platform for green plant genomics. Nucleic Acids Res 40, D1178–D1186

61. One Thousand Plant Transcriptomes Initiative (2019) One thousand plant transcriptomes and the phylogenomics of green plants. Nature 574, 679–685

62. Gu, K.J. et al. (2022) GinkgoDB: an ecological genome database for the living fossil, Ginkgo biloba. Database 2022, baac046.

63. Ronquist, F. et al. (2012). MrBayes 3.2: efficient Bayesian phylogenetic inference and model choice across a large model space. Syst Biol 61, 539–542

64. Gelman, A. & Rubin, D.B. (1992) Inference from iterative simulation using multiple sequences. Stat Sci 7, 457–472

65. Postma, M. & Goedhart, J. (2019) PlotsOfData-A web app for visualizing data together with their summaries. PLoS Biol 17, e3000202

