## Supplemental Figures for "Control of stem cell behavior by CLE–JINGASA signaling in the shoot apical meristem in *Marchantia polymorpha*"

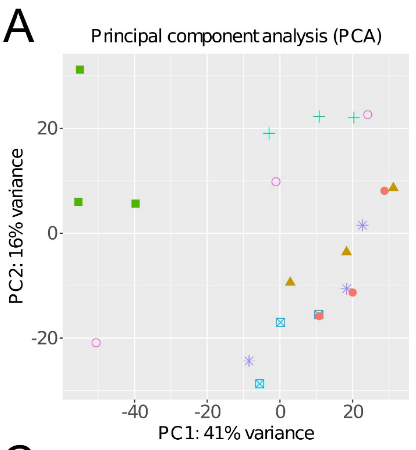


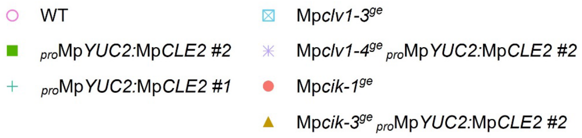


B

| | $_{pro}$Mp*YUC2*: Mp*CLE2* #2 | $_{pro}$Mp*YUC2*: Mp*CLE2* #1 | Mp*clv1-3*$^{ge}$ | Mp*clv1-4*$^{ge}$ $_{pro}$Mp*YUC2*: Mp*CLE2* #2 | Mp*cik-1*$^{ge}$ | Mp*cik-3*$^{ge}$ $_{pro}$Mp*YUC2*: Mp*CLE2* #2 |
|---|---|---|---|---|---|---|
| DEGs | 749 | 620 | 322 | 261 | 306 | 476 |
| DETFs | 21 | 19 | 20 | 14 | 12 | 10 |

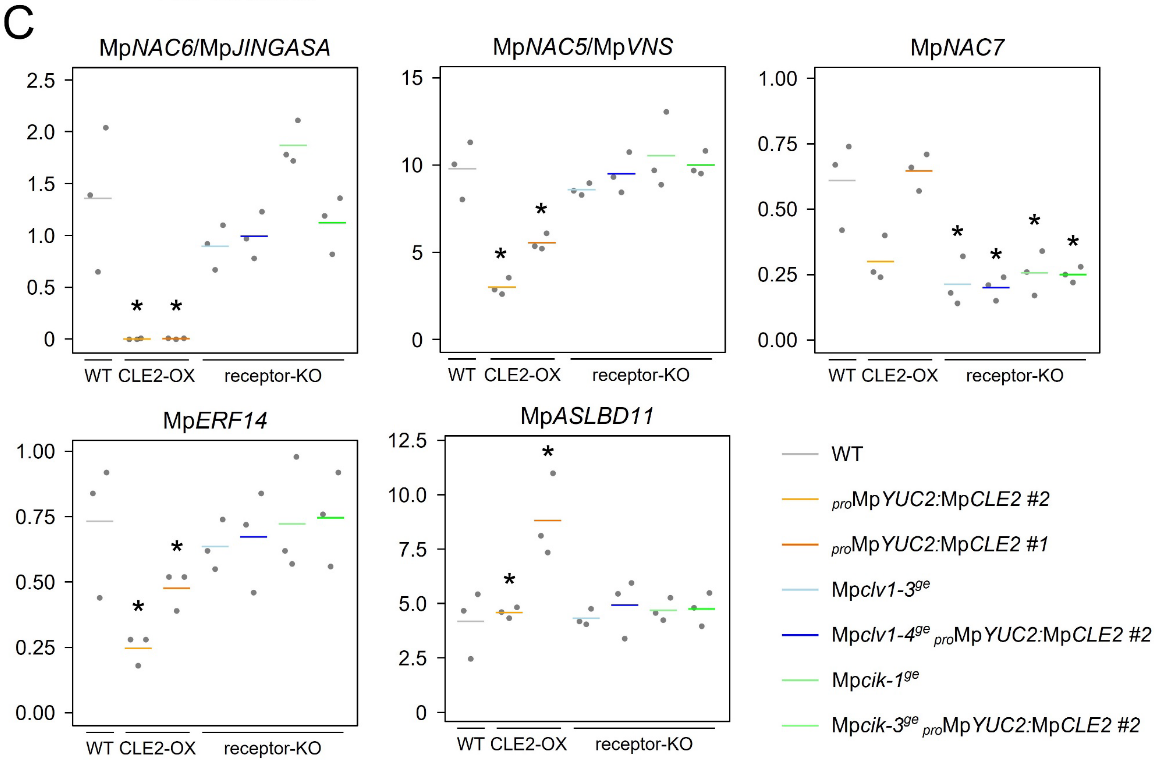


**Figure S1. RNA-seq for identification of downstream targets of MpCLE2 signaling.** (**A**) Principal component analysis (PCA) of RNA-seq data. The first and second principal component values (PC1 and PC2) of each sample are plotted. (**B**) The number of differentially expression genes (DEGs) and differentially expressed transcription factors (DETFs) in MpCLE2-OX lines and Receptor-KO alleles. (**C**) Transcripts per million (TPM) of specifically detected DETFs in MpCLE2-OX lines and receptor-KO alleles, respectively. In **C**, data is represented by mean (bars) and individual data points (dots). Asterisks indicate differential expression compared to WT (FDR<0.1).

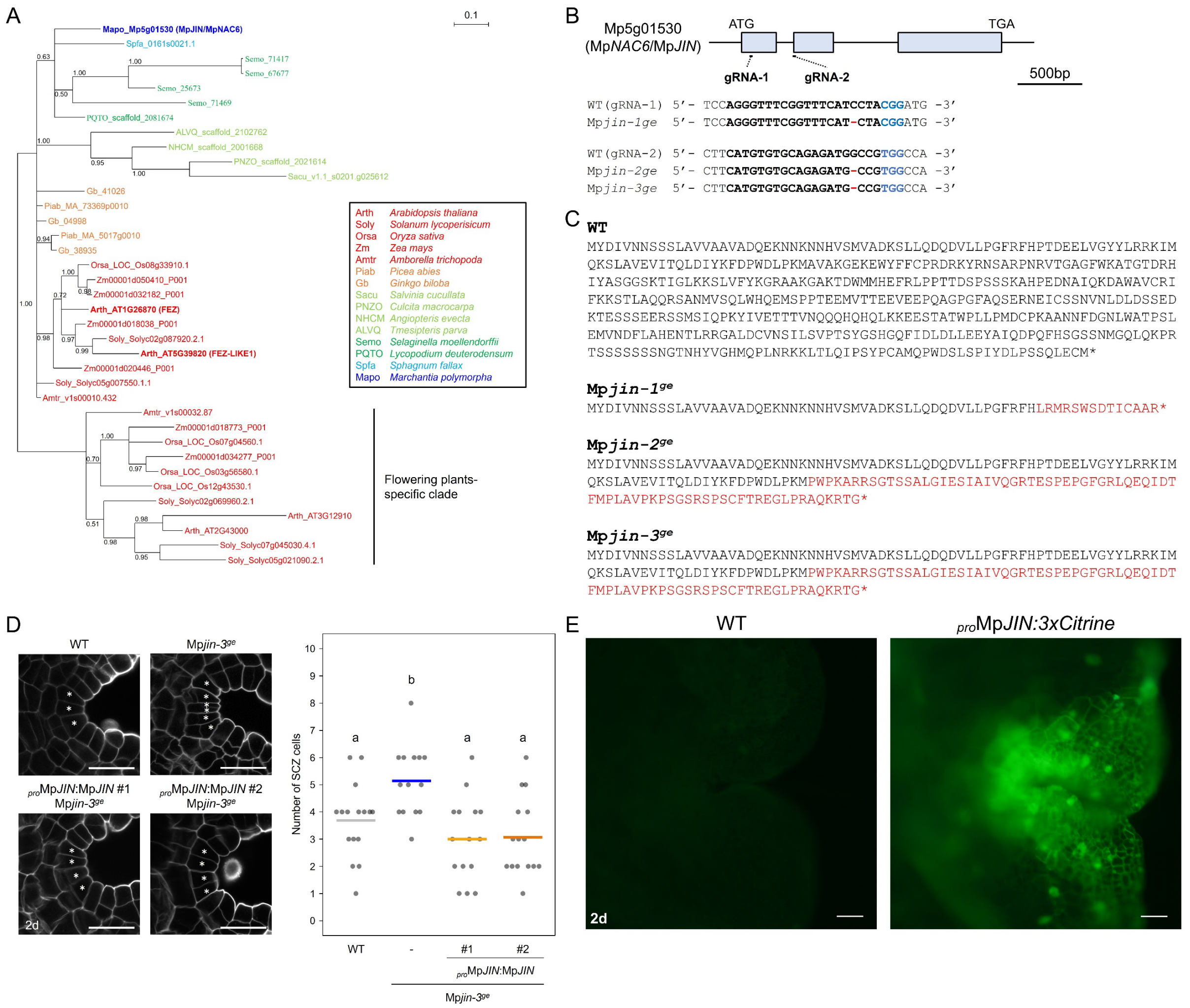


**Figure S2. Molecular genetic analysis on Mp*JIN*** (**A**) A phylogenetic tree of JIN/FEZ subfamily, generated with a Bayesian method based on the conserved NAC domain. The posterior probabilities of trees are shown at the nodes. (**B**) (top) Structure of MpJIN/Mp5g01530 locus with the position of designed guide RNA (gRNA). Exons are shown as boxes. (bottom) Genotyping of genome editing alleles. gRNA sequence is in bold, and PAM sequence is in blue. Deleted bases are indicated with hyphens in red. (**C**) WT and mutant protein sequences deduced from the genomic DNA sequences are indicated. Sequences different from wild type (WT) are in red. Asterisks indicate translational termination. (**D**) Complementation of an Mp*jin*$^{ge}$ allele by Mp*JIN* transgene. (left) Confocal imaging of the SCZ in 2-day-old gemmalings. Asterisks indicate the SCZ cells. Scale bars represent 25 µm. (right) Quantification of the cell number in the SCZ (n=14-16). Data is represented by mean (bars) and individual data points (dots). Two-way ANOVA with Tukey’s post hoc test. Means sharing the superscripts are not significantly different from each other, $p < 0.05$. (**E**) Fluorescence imaging of Mp*JIN* promoter activity at the apical notches of 2-day-old gemmalings. (left) Wild type (Tak-1) plant. (right) $_{pro}$Mp*JIN:3xCitrine* plant. Scale bars represent 100 µm.

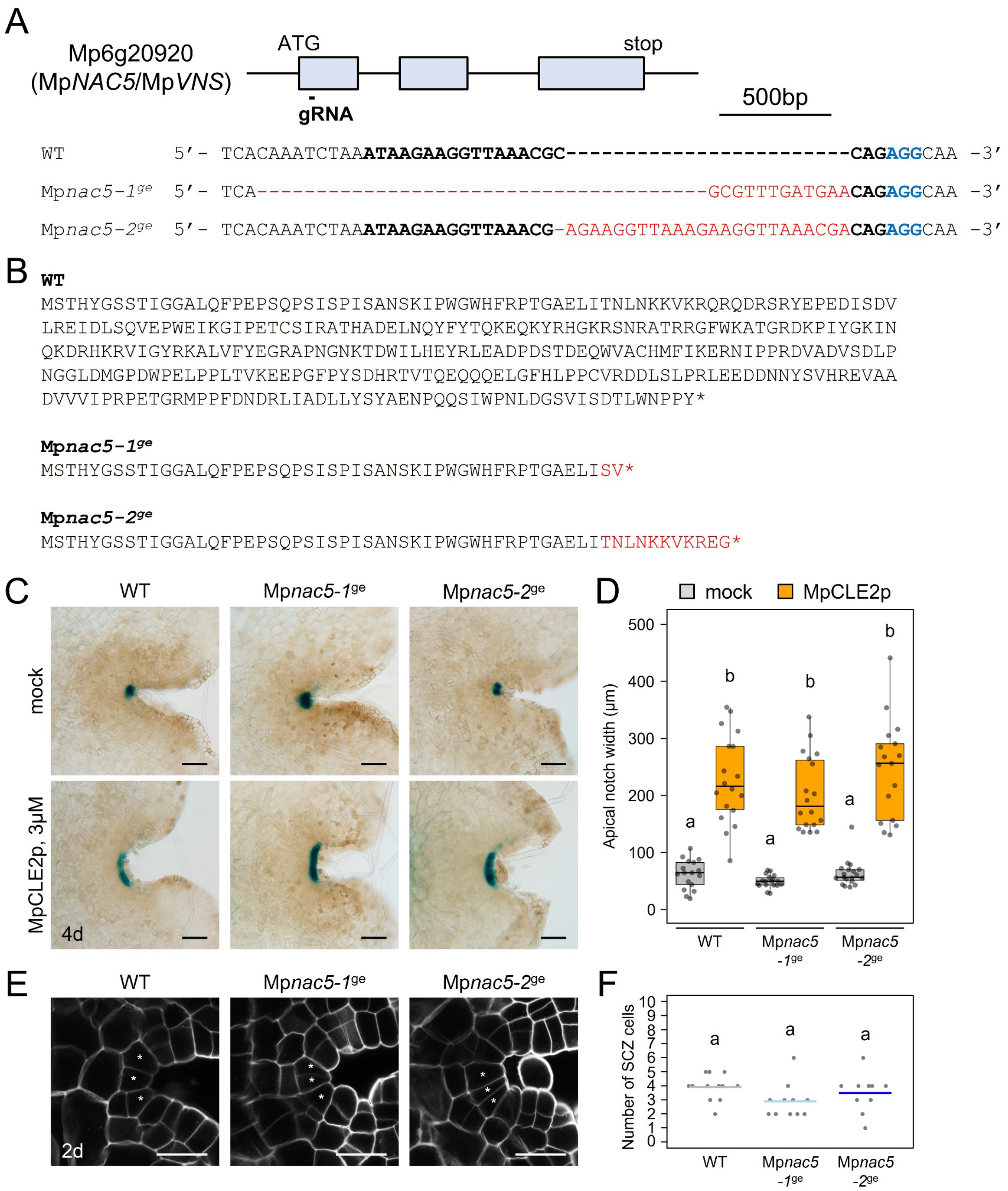


**Figure S3. Genome editing of Mp*NAC5*/Mp*VNS*.** (**A**) (top) Structure of Mp6g20920/MpNAC5/MpVNS locus with the position of designed guide RNA (gRNA). Exons are shown as boxes. (bottom) Genotyping of Mp*nac5*$^{ge}$ alleles. gRNA sequence is in bold, PAM sequence is in blue, and insertion/deletion is in red. (**B**) WT and mutant proteins deduced from the genomic DNA sequences. Mutant protein sequences different from WT are indicated in red. Asterisks indicate translational termination. (**C**) $_{pro}$Mp*YUC2:GUS* marker in 4-day-old gemmalings grown with or without 3 μM MpCLE2 peptides. Genotypes of MpNAC5 are indicated above the panels. (**D**) Quantification of the apical notch width (n=17-19). (**E**) Confocal imaging of the SCZ in 2-day-old gemmalings. Asterisks indicate the SCZ cells. (**F**) Quantification of the cell number in the SCZ (n=10-12). In **D**, the boxes show the median and interquartile range, and the whiskers show the 1.5x interquartile range. Individual data points are plotted as dots. In **F**, data is represented by mean (bars) and individual data points (dots). Two-way ANOVA with Tukey's post hoc test in **C**, **E**, **G** and **I**. Means sharing the superscripts are not significantly different from each other, $p < 0.05$. Scale bars represent 100 μm (**C**) or 25 μm (**E**).

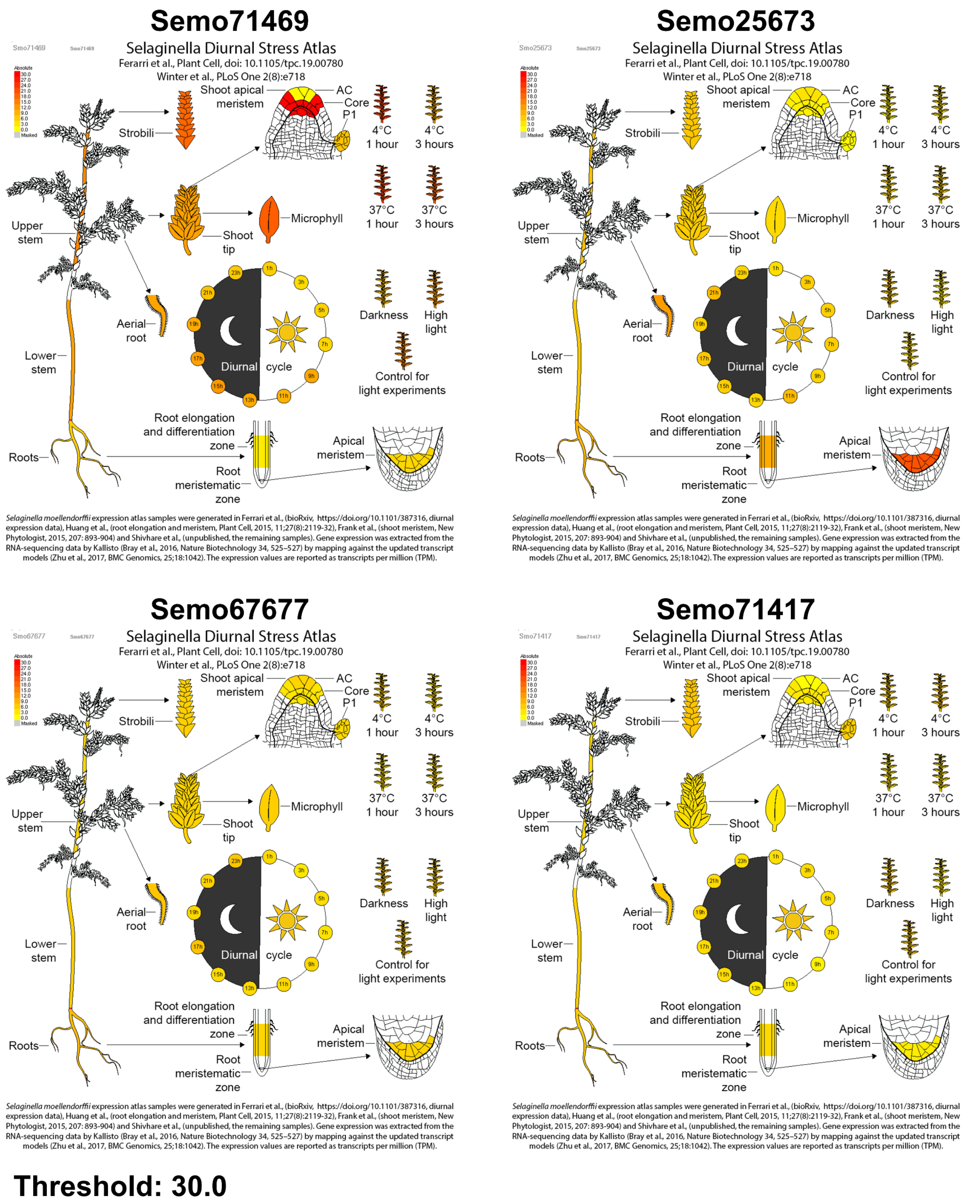


**Figure S4. Expression of JIN/FEZ homologs in *Selaginella moellendorffii* gene expression atlas.** Expression levels of JIN/FEZ homologs were obtained from the gene expression atlas using the eFP Browser website (http://bar.utoronto.ca/efp_selaginella/cgi-bin/efpWeb.cgi)[44]. Signal threshold values for visualization are set as 30.0 for comparison among genes.
